# Bearing the brunt: Mongolian khulan (*Equus hemionus hemionus*) are exposed to multiple influenza A strains

**DOI:** 10.1101/357905

**Authors:** Eirini S. Soilemetzidou, Erwin de Bruin, Gábor Á. Czirják, Bayarbaatar Buuveibaatar, Petra Kaczensky, Marion Koopmans, Chris Walzer, Alex D. Greenwood

## Abstract

The majority of influenza A virus strains are hosted in nature by several Anseriformes and Charadriformes birds. A minority of strains have been able to cross species boundaries and establish themselves in novel non-avian hosts. Influenza viruses of horses, donkeys, and mules represent successful cases of avian to mammal influenza virus adaptation. Mongolia has over 3 million domestic horses and is home to two wild equids, the Asiatic wild ass (*Equus hemionus hemionus*), and Przewalski horse (*Equus ferus przewalskii)*. Domestic and wild equids are sympatric across most of their range in Mongolia. Epizootic influenza A virus outbreaks among Mongolian domestic horses have been frequently recorded. However, the exposure, circulation and relation to domestic horse influenza A virus outbreaks among wild equids is unknown. We evaluated serum samples of Asiatic wild asses in Mongolia for antibodies against influenza A viruses, using a serological assay. We detected antibodies against hemagglutinin (H) H1, H3, H5, H7, H8 and H10 influenza A viruses. Asiatic wild asses may represent a previously unidentified influenza A virus reservoir in an ecosystem shared with populations of domestic horses in which influenza strains circulate.

## Importance

Influenza A virus monitoring of domestic animals is often undertaken independent of the ecological context in which the animals exist. Domestic horses in Mongolia are sympatric and have largely unconstrained contact with wild equids within the Mongolian steppe ecosystem. Our results suggest that the Asiatic wild asses are exposed to known equine influenza A strains H3N8 and H7N7 and additional influenza strains not previously described in horses. H7N7 has been considered to be absent from the region but is clearly not. Asiatic wild asses may therefore represent an influenza A reservoir of relevance to the domestic and wild equid population of eastern Asia.

## Introduction

Historically, two major strains of Equine Influenza virus (EIV) have caused influenza virus outbreaks in domestic equids. The first identified EIV, influenza A/H7N7 or equine-1, was isolated from horses in 1956 (1). Influenza A/H3N8 or equine-2 was subsequently reported and remains the major cause of equine influenza (*2*). While H7N7 EIV is thought to be equine-specific with limited but unique variation in the HA gene (*3*), H3N8 EIV appear to bind to avian-like receptors in the upper respiratory tract of horses suggesting a recent avian origin of the strain (*4*). Moreover, previous H3N8 influenza virus outbreaks in dogs (*5*), their isolation from a Bactrian camel in Mongolia (*6*), and some evidence for human infection (*7*), indicate that horses are not the only host for H3N8 viruses, and that their zoonotic potential might be underappreciated. Mongolia, with a current population of domestic horses exceeding 2 million, has suffered several EIV outbreaks (8). The first two outbreaks, 1974-75 and 1983-84 (*9*), were caused by H7N7 EIV and the last three, 1993-94, 2007-08 (*9*) and 2011 were caused by H3N8 EIV. After 1984, H7N7 EIV was not isolated and is considered extinct in the region.

In addition to domestic horses, Mongolia is home to the Przewalski horse (*Equus ferus przewalskii)*, and hosts the biggest population of Asiatic wild ass (or Khulan, *Equus hemionus hemionus*) in Central Asia (10). The distribution of khulan overlaps with other free-living ungulate species, such as goitered gazelles (*Gazella subgutturosa*), Mongolian gazelles (*Procapra gutturosa*), and wild Bactrian camels (*Camelus ferus*). Most importantly their distribution overlaps with local livestock including domestic horses which outnumber wild ungulates by several orders of magnitude. Disease transmission between domestic and free-living populations is possible through sharing pasture and waterholes. EIV outbreak dynamics in wild equids from Central Asia are poorly understood. In 2007 an H3N8 influenza (A/equine/Xinjiang/4/2007) outbreak was reported in a Przewalski’s horse population in the Chinese part of the Gobi with a 5% mortality rate (11). Influenza exposure in khulans, however, remains uncharacterized. Mongolia also has a high diversity of wild birds, including migratory waterbirds, that use Mongolia as a stop-over during their annual migrations. The Central and the East Asian flyways passing through Mongolia are critical to influenza ecology (Figure 1). Therefore, we sought to investigate history of exposure to influenza viruses in wild equids, as a first step in understanding their role in the ecology of equine influenza.

**Figure 1:**
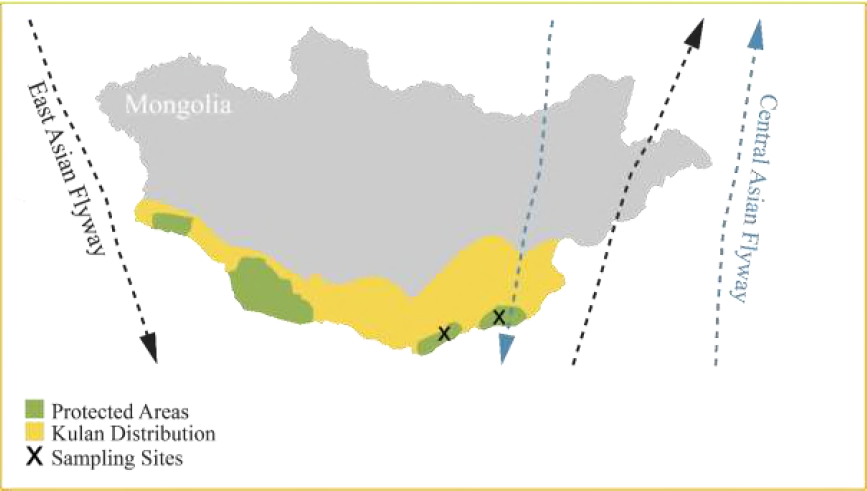
Migratory bird flyways through Mongolia. The map illustrates the relation between khulan distribution, protected areas, sampling sites and the major migratory flyways in Mongolia. The arrows represent the outside border of each migratory flyway.

## Results

### Field work

Sampling of Khulan was done in Southern Gobi, Mongolia after general anesthesia. The method of choice for efficient chemical capture in the Gobi desert is from a moving jeep. After successful detection of khulan in the steppe, and initiation of the chase, there is a cut off time of 15 min for animal welfare reasons which determines when capturing will end. Subsequently, a new khulan group needs to be found before continuing. The time to capture (from detection and initiation of the chase to reversal of the anesthesia to being ready to resume the search for a new animal) for individual animals ranged from approximately 1 hour to several days. In total, we captured 21 adult khulan (8 stallions and 13 mares). After transportation to the laboratory, blood samples from 13 animals were found to be suitable for analysis.

### Influenza A virus testing

Viral detection was attempted from nasal swabs using qPCR but no virus could be detected. The result is not surprising as 460 domestic free-ranging Bactrian camels were similarly screened yielding a single influenza A virus positive individual (6). Considering none of the animals displayed clinical symptoms of infection, the lack of actively shedding individuals is consistent with expectations.

### Influenza A virus serology

To detect exposure to influenza in non-shedding individuals, a protein microarray (PA) technique testing 32 hemagglutinin recombinant proteins (HA1-part) from type H1 to H16, as described previously (*12-14*), was used to profile the antibodies to influenza viruses in the khulan serum (Table 1). Six animals were negative, whereas 7 animals had reactivity detectable by microarray to one or more antigens. These were low levels of reactivity to H5 (2 animals), H8 and H10 antigen (1 animal each), and low to moderate titers against H1 (1 animal), and H7 (2 animals). Five khulans showed reactivity to H3-08, which is the horse influenza strain known to circulate in Mongolia. This reactivity was specific for the EIV H3 antigen, other antigens (representing strains isolated from humans) were negative (Figure 2).

**Figure 2:**
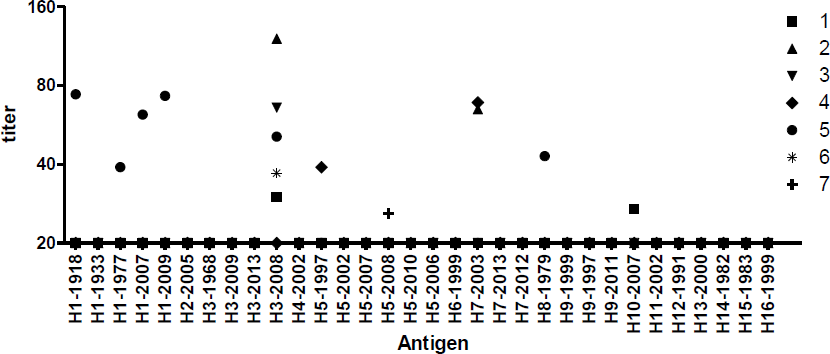
Antibody profiles in sera from khulans, expressed as titers (Y axis) of IgG reactivity to a range of influenza A HA1 antigens (X axis). The legend shows the animal number

**Table 1.** Recombinant HA1-proteins included in the protein microarray.

| **CODE** | **SUBTYPE** | **STRAIN** |
| --- | --- | --- |
| **H1-1918** | H1N1 | A/South Carolina/1/18 |
| **H1-1933** | H1N1 | A/WS/33 |
| **H1-1977** | H1N1 | A/USSR/92/1977 |
| **H1-2007** | H1N1 | A/Brisbane/59/2007 |
| **H1-2009** | H1N1 | A/California/6/2009 |
| **H2-2005** | H2N2 | A/Canada/720/05 |
| **H3-1968** | H3N2 | A/Aichi/2/1968(H3N2) |
| **H3-2009** | H3N9 | A/VICTORIA/210/2009 |
| **H3-2013** | H3N2 | A/Switzerland/9715293/2013 |
| **H3-2008** | H3N8 | A/equine/Gansu/7/2008 |
| **H4-2002** | H4N6 | A/mallard/Ohio/657/2002 |
| **H5-2997** | H5N1 | A/Hong Kong/156/97 |
| **H5-2002** | H5N8 | A/duck/NY/191255-59/2002(H5N8) LP |
| **H5-2007** | H5N3 | A/duck/Hokkaido/167/2007 |
| **H5-2008** | H5N1 | A/chicken/Egypt/0879-NLQP/2008 |
| **H5-2010** | H5N1 | A/Hubei/1/2010 |
| **H5-2006** | H5N1 | A/Turkey/15/2006 (clade 2.2) |
| **H6-1999** | H6N1 | A/quail/HK/1721-30/99 |
| **H7-2003** | H7N7 | A/Chicken/Netherlands/1/03 |
| **H7-2013** | H7N9 | A/chicken/Anhui/1/2013 |
| **H7-2012** | H7N3 | A/chicken/Jalisco/CPA1/2012 |
| **H8-1979** | H8N4 | A/pintail duck/Alberta/114/1979 |
| **H9-1999** | H9N2 | A/Guinea fowl/Hong Kong/WF10/99 |
| **H9-1997** | H9N2 | A/chicken/Hong Kong/G9/97 (G9 lineage) |
| **H9-2011** | H9N2 | HA1 (H9N2) A/Chicken/India/IVRI-0011/2011 |
| **H10-2007** | H10N7 | A/blue-winged teal/Louisiana/Sg00073/07 |
| **H11-2002** | H11N2 | A/duck/Yangzhou/906/2002 |
| **H12-1991** | H12N5 | A/green-winged teal/ALB/199/1991 |
| **H13-2000** | H13N8 | A/black-headed gull/Netherlands/1/00 |
| **H14-1982** | H14N5 | A/mallard/Astrakhan/263/1982new |
| **H15-1983** | H15N8 | A/duck/AUS/341/1983 |
| **H16-1999** | H16N3 | A/black-headed gull/Sweden/5/99 |

The two khulan serum samples which reacted with H7 antigen, reacted specifically to the Dutch H7N7 strain (H7-03, A/Chicken/Netherlands/1/03), but not to the Chinese H7N9.

We cannot exclude that multiple known or unknown strains of H1, H5, H8 and H10 cross reacted due to the viscous nature of khulan serum or that the viral strains eliciting the immune response are divergent from known strains. Confirmation of the results using hemagglutination inhibition (HI) assays was not possible because the serum was severely hemolysed and a-specific agglutination was detected in the control well without virus. Therefore, the specific influenza viruses responsible for eliciting the development of these antibodies remain unknown.

## Discussion

Our results suggest that wild khulan in Mongolia’s Southern Gobi are exposed to multiple influenza A viruses, including horse H3N8. Although we could not detect viral genomes to further define the equine atypical strains our results suggest equids may be susceptible to more influenza viruses than previously considered. Virus detection is often limited by the short window in which the virus is present, and therefore screening for antibodies, which often persist longer than the virus itself, provides information about past infections and virus diversity in animal populations (14). While sampling of twenty-one individual animals may seem low, one has to keep in mind that khulans are extremely skittish animals, and normally flee human presence even when separated by several kilometers distance. Anesthesia and sampling of non-domestic equids, particularly under the physically challenging and remote environment of the Gobi Desert, can be difficult, for both animals and humans, and not always successful (15). Furthermore, khulan are a red list species globally and nationally and capture permits are granted only after careful evaluation of the risks and benefits. On these grounds, capture permits for the mere sampling of an endangered species without an imminent need have little chance of approval.

The difficult terrain, with dry river beds, low mountains, bushes, shrubs and desert basins, severely restricts successful outcomes. Capture (from detection and initiation of the chase to reversal of the anesthesia to being ready to resume the search for a new animal) for one individual takes approximately 1 hour under the best conditions but ranges to several days, if khulan are not found in the vast Gobi ecosystem (16). In our study, the numbers of animals captured exceeded the expectations for our short 2 week window. These challenges need to be taken into account when evaluating this study.

We found evidence for exposure to influenza viruses with a hemagglutinin of subtype H7. H7N7 equine influenza is considered extinct in the region. In China, an outbreak of avian H7N9 influenza virus has been in progress since 2013, but the protein microarray results suggest the virus eliciting the immune response was different from current H7N9 strain. Other H7 subtypes circulate in wild birds in Southeast Asia in addition to H7N9, and further research should clarify to which specific H7 influenza virus khulan might be exposed in Mongolia. Khulans may have been infected with an H1 strain during an H1N1 pandemic in 2009 (*17*), and there is evidence that donkeys, a domestic equid, are susceptible to highly pathogenic H5N1 influenza strains (*18*). H8 has been described in American mink (*19*), while H10 is a known human pathogenic strain linked to direct poultry contact and H10 strains can also infect mammals such as harbor seals (*20*).

The most commonly antibodies were against H3 EIV HA1 antigens, consistent with data on circulation of these viruses in horses. Individual animals tested positive for HA1 of influenza A H1, H5, H7, H8 and H10 viruses, suggesting that sporadic infections with viruses belonging to these subtypes have occurred. A possibility is that these viruses co-circulate with H3N8 among equids in Central Asia, but occasional introduction from exposure to wild birds or their droppings is a possible alternative. The upper respiratory tract of the horse express sialic acid 2,3-linked receptors, which are similar to those in wild aquatic birds. Because of this similarity in avian and equine respiratory biology (*21*) it is possible that equids are susceptible to a broader spectrum of influenzas than other mammals.

Although susceptibility of wild equids to new influenza strains may not pose a threat to their conservation status, it might represent an overlooked ecological niche for influenza virus and an alternative route of infection for other wild and domestic animals. Further epidemiological investigation of wild equids from Central Asia should clarify the diversity of influenza virus strains that infect wild equids and help to establish the monitoring of transmission of influenza virus between wild and domestic equids in the area.

## Materials and Methods

The study took place in the Southern Gobi Desert in Mongolia, and was approved by the ethical committee of the University of Veterinary Science in Vienna (ETK-15/03/2016) and the Mongolian Government (05/5656). Twenty-one adult khulan (8 stallions and 13 mares) were anesthetized and nasal swabs and blood samples collected. The animal sampling expedition, was part of a radio collaring project, in which habitat fragmentation, due to new mining-related infrastructures in Southern Gobi were investigated. Khulans were captured in two different locations, one near the mining-infrastructure site and one near Ergeliin Zoo protected area. All animals were darted from a moving jeep, using a Daninject JM CO2 darting gun (Walzer et al. 2006). None of the khulan demonstrated clinical symptoms of EIV or other infectious diseases when handled. Samples were stored immediately at -20°C in a portable freezer in Mongolia, transported on dry ice to Austria in full compliance with the Convention on International Trade in Endangered Species (CITES) and stored at −80°C until laboratory analysis at the Research Institute of Wildlife Ecology, University of Veterinary Medicine, Vienna. Due to field conditions and the absence of a mobile laboratory, blood samples couldn’t be proceeded on site, so that only 13 of 21 serum samples could be processed and all were severely hemolysed.

200 μl transport medium from the swabs were extracted using the RTP DNA-RNA virus mini kit (Stratec Molecular GmbH, Germany). The RNA was reverse transcribed cDNA using an influenza specific primer followed by PCR amplification. All the reactions were performed in a single tube, by using the SensiFast^TM^ Probe Lo-ROX One-Step kit (Bioline). Negative and positive controls were included in all experiments. Reactions were carried out in 96-well plates. Influenza M genes primers and probes used were as follows: Primer INF FW 5’- AGA TGA GYC TTC TAA CCG AGG TCG -3’, Primer INF RV 5’- TGC AAA NAC ATC YTC AAG TCT CTG -3’, Probe INF 6 FAM- TCA GGC CCC CTC AAA GCC GA –TAMRA. qPCR was performed using an ABI 7500 machine, p with the cycling conditions: 10 min at 48°C (RT reaction), 2 min at 95°C, 40 cycles of 3 sec at 95°C and 30sec at 60°C and held for 1 min at 60°C for the data collection.

A protein microarray (PA) technique as described previously *(9, 10, 11)*, was used to identify the influenza virus strains in the kulan serum samples. Samples where inactivated in a water bath at 56° C for 4 hours due to regulations for testing of animal samples from food and mouth disease endemic regions. Serum samples from 3 kulans were tested against different secondary antibodies in order to determinate the highest sensitivity; protein A, protein G and anti-Horse. Anti-horse IgG showed highest overall response.

Briefly, thirty-two recombinant proteins of different influenza A virus antigens were printed onto 16-pad nitrocellulose Film-slides (Oncyte avid, Grace Bio-labs, Bend, OR, USA). All presently known influenza A virus HA-types are represented on the array (Table 1), except for hemagglutinin type 17 and 18, as those are only detected in bats. Slides were treaded with Blotto-blocking buffer to avoid non-specific binding (Thermo Fischer Scientific Inc., Rockford, MA, USA) for 1 hour at 37°C in a moist chamber. After washing the slides were incubated with a fourfold dilution series of the kulan serum starting from 1:20 to 1:1280. After 1 hour incubation at 37 °C, slides were washed and incubated with a 1:500 dilution of the anti-horse IgG conjugated to Alexafluor 647 (Jackson immunoresearch). A last washing step was done to remove unbound conjugate, after which the slides were dried and scanned using a Powerscanner (Tecan). Spot intensities were determined, and titer heights were calculated by curve fitting using R (R Statistical Computing, version 3.1.0, Vienna, Austria). Titers less than 20 were set to 20.

## Acknowledgments

We acknowledge the FUW-Advanced Design Studio for their insightful suggestions for the map illustration. We thank the Ministry of Nature, Environment and Tourism of Mongolia, Dashzeveg Tserendeleg, Otgonsuren Avirmed, Enkhtuvshin Shiilegdamba (WCS), Nyamdorj Barnuud (SEA), Dennis Hosack, and Purevsuren Tsolmonjav (OT) for the logistical and practical support during khulan capture.

